# A Comprehensive Study of *Piper chaba* (Chui jhal): Phytochemistry and Antioxidant Effects

**DOI:** 10.1101/2024.12.15.628592

**Authors:** Monika Kundu, Konika Kundu, Farhana Akter Joty

## Abstract

*Piper chaba*, a valuable medicinal plant from the family Piperaceae is widely cultivated for its use in traditional medicine and as a spice. This study aimed to evaluate the phytochemical composition and antioxidant potential of *Piper chaba*. Phytochemical analysis was conducted using standard chemical reaction methods, and antioxidant activity was assessed through the free radical scavenging assay. The phytochemical screening revealed the presence of alkaloids, carbohydrates, flavonoids, glycosides, resin, steroids, terpenoids, and tannin in *Piper chaba*. The presence of flavonoids indicates significant antioxidant potential. By using the DPPH scavenging method, the plant showed an IC_50_ value of 5.81. Further analysis is required to identify the potentiality of the plant in various areas.

## 1. Introduction

For many years, medicinal plants have been essential to human health and wellbeing since they are the main source of bioactive substances used to treat a wide range of illnesses. Because plants include phytochemicals like alkaloids, flavonoids, tannins, and glycosides. Traditional medicine, which is practiced in many different cultures, frequently depends on them for its medicinal benefits. The pharmacological activities of plants, such as their antibacterial, anti-inflammatory, and antioxidant properties, are greatly influenced by these substances (Roy, 2022).

The family Piperaceae, which comprises several plants well-known for their culinary and therapeutic use, includes *Piper chaba*. It is a native of South and Southeast Asia. It is frequently used as a spice and is prized in traditional medical systems for its capacity to reduce pain, inflammation, and microbial infections. It is also known locally as “Chui jhal” and is especially well-liked in places like Bangladesh and India, where its stems and roots are utilized in a variety of recipes (Salehi, 2019).

Chronic diseases such as cancer, neurological diseases, and cardiovascular disorders are largely caused by oxidative stress, which is brought on by an imbalance between the body’s antioxidants and free radicals (Pizzino, 2017). Plant-based natural antioxidants have drawn a lot of interest as possible treatment options for oxidative damage. Often present in medicinal plants, flavonoids are a class of phytochemicals known for their strong antioxidant and free radical-scavenging capabilities. Despite the long history of traditional use, there is still little thorough research examining the phytochemical composition and antioxidant potential of *Piper chaba*(Panche, 2016). According to preliminary data, the plant may contain beneficial bioactive substances including flavonoids that support its therapeutic qualities. This study aims to assess This plant’s phytochemical profile and antioxidant activity in order to better comprehend its medicinal potential. These revelations might open up the possibilities for its incorporation into contemporary pharmaceutical applications (Ashokkumar, 2021).

### Identification & Collection of plant material

A healthy plant was selected which means the plant chosen had no disease. The plant chui jhal (*Piper chaba)* was collected from Jashore district, village of monirampur area of Bangladesh on December 24, 2017, and this plant was implanted on 26 December 2017 in Noakhali Science and Technology University, Sonapur, Noakhali. Then small pieces of plant material were collected from Jashore district for the Phytochemical and Antioxidant test. Then wash the plant material to remove dust.

## 2. Methods

### Extraction procedure Drying and Grinding

The plant’s stem and leaves were initially separated. After that, the gathered plant material was cleaned with water and segregated from other unwanted materials. The well cleaned plant material was chopped into tiny pieces and allowed to dry for three days in a moderate amount of sunshine. Because it lowers the water content of the plant material, the fully dried plant materials remained dry in the oven for 45 to 50 minutes at 45_o_C. Water is known to promote microbial growth. A suitable grinder was then used to grind these into a coarse powder. Until analysis began, the powder was maintained in a dry, dark, and cool environment in an airtight container. After that, the powder was employed extensively in the study.

### Cold Extraction

The cold extraction procedure was used to remove the components. In a glass container, 50g of powdered powder was steeped in 500ml of ethanol for 12–15 days. This container was covered with parafilm paper and maintained in a dark location. Every two to three hours, it shakes for cold extraction. The extract was then filtered through a piece of filter paper to remove the plant debris. Fresh plant extract was discovered for this procedure. After that, it uses a spatula to remove a container and stored it in the fridges.

### Phytochemical test

Phytochemical examinations were carried out for the extract as per the standard methods (Nortjie, 2022).

### Antioxidant test

#### DPPH solution preparation

To create a DPPH solution with a concentration of 40 μg/mL, 4 mg of DPPH powder was weighed and dissolved in 100 mL of ethanol. The amber reagent bottle was used to make the solution, which was then stored for forty-five minutes in a dark, ice-filled, lightproof box.

### Assay Procedure

To obtain the parent solution (concentration 1000 μg/mL), calculated amounts of various extractives (about 2 mg) were measured and dissolved in ethanol.Serial concentration dilution was 500 μg, 100 μg, 50 μg, 10 μg, 5 μg, 1 μg. To create the concentration six volumetric flasks were taken and test tubes were wrapped in foil paper.

First,1mL of stock solution was mixed with 19mL of solvent (ethanol) to get a 500 μg concentration. Then, to create a 100μg concentration, 1 ml of the 500μg concentration solution was combined with 4 ml of ethanol solvent. Once more, 1 milliliter of 500 μg concentration was combined with 9 milliliters of ethanol solvent to create a 50 μg concentration. Combine 1 ml of 50 μg concentration with 9 ml of solvent (ethanol) to get a 10 μg concentration. Then, 9ml of solvent (ethanol) was combined with 1ml of 50μg concentration. Additionally, a 1μg concentration was made by combining 4ml of solvent (ethanol) with 1ml of 5μg concentration. Finally, a blank solution was made by combining 3mL of DPPH solution with 1 milliliter of solvent (ethanol).

The extract solution and DPPH solvent are then mixed in varying concentrations (1:3), meaning that 1mL of extract solution and 3mL of DPPH solution are added. After 45 minutes of incubation at room temperature (25º C) in a dark environment, the absorbance at 517 nm was measured using a UV spectrophotometer against black ethanol.

## 3. Results & Discussion

### Phytochemical test

Phytochemical test:
| Test | Result |
| --- | --- |
| Steroids | + |
| Alkaloids | + |
| Carbohydrates | + |
| Flavonoids | + |
| Glycosides | + |
| Resins | + |
| Tannins | + |
| Terpenoids | + |

Here,

+ = Presence

#### Ascorbic Acid

DPPH radical scavenging activity of Ascorbic acid (standard)

**Figure 1:**
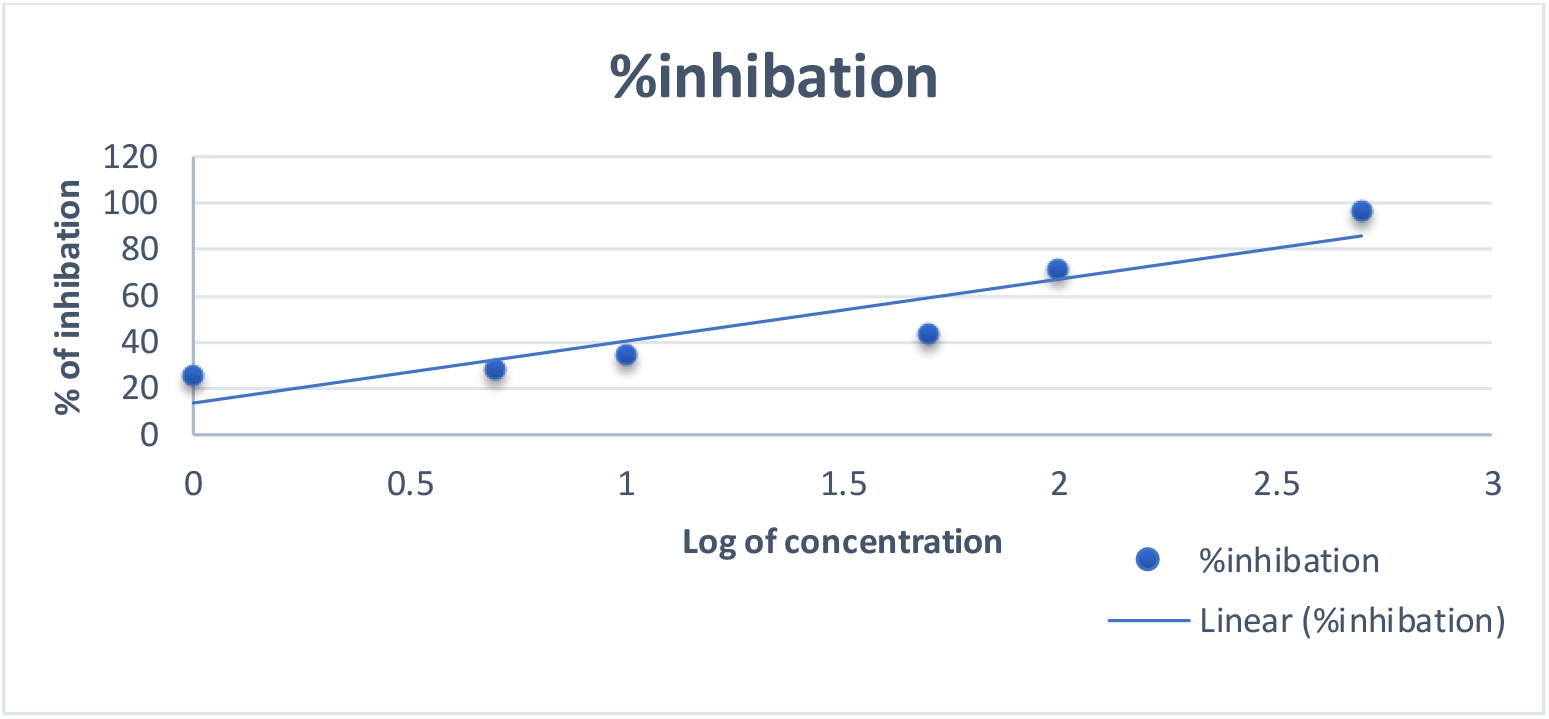
Log of Concentration vs. % Inhibition graph of ascorbic acid for evaluating IC_50_. Blank absorbance = 1.33 and IC_50_= 1.6

##### Piper chaba

DPPH radical scavenging activity of *Piper chaba*

**Figure 2:**
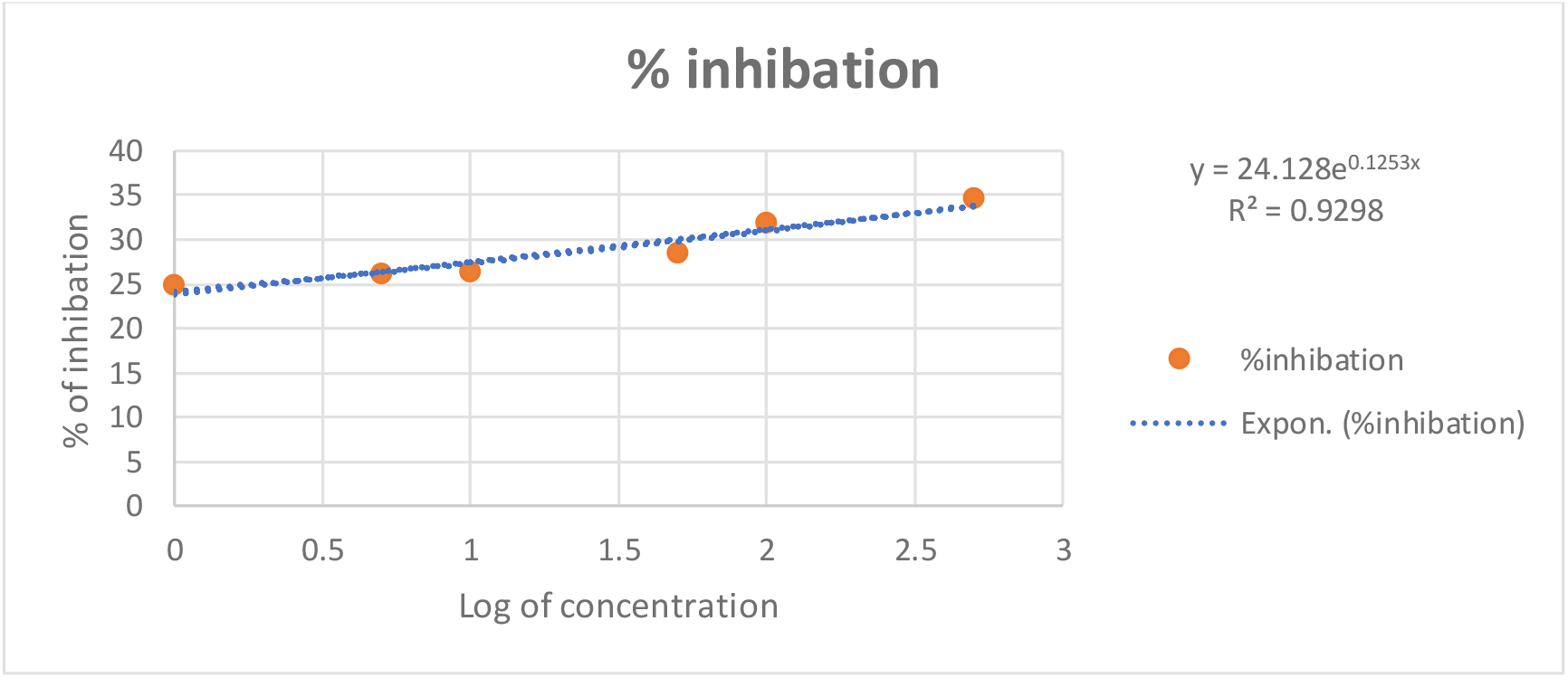
Log of Concentration vs. % Inhibition graph for evaluating IC_50_. IC_50_= 5.81

The phytochemical test of *Piper chaba* indicated the presence of Steroids, Alkaloids, carbohydrates, flavonoids, glycosides, resin, tannin, and terpenoids.

IC_50_ is the concentration at which 50% of the total DPPH free radical is scavenged and neutralized. Here compare the IC_50_ value between ascorbic acid and *Piper chaba*. The lowest IC_50_ value indicates the highest antioxidant activity. Using the exponential (y=mx) equation on this graph for y=50 value and determining the value of x. This x value indicates the IC_50_ value. Here the IC_50_ value of *Piper chaba* is 5.81μg/ml and also get IC_50_ value of ascorbic acid is 1.6μg/ml. This value indicates that piperine shows good antioxidant activity.

### Conclusion

*Piper chaba* belongs to the family of Piperaceae and genus Piper and is locally known as chui jhal. The Piperaceous plants contain a wide range of chemical and unique pharmacologically active compounds and different parts of this plant are being used for the treatment of different diseases. From this study, it can be concluded that the species is effective in scavenging free radicals and has the potential to be a powerful antioxidant. The extract of Piper chaba was compared with the ascorbic acid and determined the IC_50_ by using a UV spectrophotometer. A lower IC_50_ value indicates higher antioxidant activity. This value indicates that the plant shows good antioxidant activity and can be explored further to determine various pharmacological activities.

## Notes

### Competing Interest Statement

The authors have declared no competing interest.

